# Efficient pan-cancer whole-slide image classification and outlier detection using convolutional neural networks

**DOI:** 10.1101/633123

**Authors:** Seda Bilaloglu, Joyce Wu, Eduardo Fierro, Raul Delgado Sanchez, Paolo Santiago Ocampo, Narges Razavian, Nicolas Coudray, Aristotelis Tsirigos

## Abstract

Visual analysis of solid tissue mounted on glass slides is currently the primary method used by pathologists for determining the stage, type and subtypes of cancer. Although whole slide images are usually large (10s to 100s thousands pixels wide), an exhaustive though time-consuming assessment is necessary to reduce the risk of misdiagnosis. In an effort to address the many diagnostic challenges faced by trained experts, recent research has been focused on developing automatic prediction systems for this multi-class classification problem. Typically, complex convolutional neural network (CNN) architectures, such as Google’s Inception, are used to tackle this problem. Here, we introduce a greatly simplified CNN architecture, PathCNN, which allows for more efficient use of computational resources and better classification performance. Using this improved architecture, we trained simultaneously on whole-slide images from multiple tumor sites and corresponding non-neoplastic tissue. Dimensionality reduction analysis of the weights of the last layer of the network capture groups of images that faithfully represent the different types of cancer, highlighting at the same time differences in staining and capturing outliers, artifacts and misclassification errors. Our code is available online at: https://github.com/sedab/PathCNN.

## Introduction

Examination of hematoxylin & Eosin (H&E) images of surgical resections or tissue biopsies by pathologists is often the first step in identifying and characterizing a tumor. Given the size of such images (from tens to hundreds of thousands of pixels wide), the complexity and heterogeneity of neoplastic and non-neoplastic pathology, this task is non-trivial with some risk to miss small tumor foci or misclassifying minor variants or features present in the image. The benefit of distinguishing non-neoplastic conditions from neoplastic tissue is obvious, but quickly and accurately differentiating between subtypes can also be critical for initiating targeted therapies [1]. For example, in the case of lung cancer, the efficacy of conventional chemotherapies differs for the two major subtypes of non-small cell lung cancer (NSCLC), squamous cell carcinoma and adenocarcinoma. Certain agents may be less efficient [2] or more invasive for patients presenting squamous cell carcinoma [3]. On the other hand, adenocarcinoma is more likely to contain genetic mutations which may be treated with targeted therapy, such as EGFR mutations or ALK rearrangements [4]. Although neoplastic subtypes are often visually distinct, diagnostic agreement for classifying adenocarcinomas and squamous cell carcinomas of the lung has been found to be relatively low (κ = 0.41 - 0.46 among community pathologists, κ = 0.64 - 0.69 among pulmonary pathology experts and κ = 0.55 - 0.59 among all pathologists under study) [5]. The difference between community pathologists and pulmonary pathology experts suggests that automated technology may aid the diagnostic process in areas lacking specialized expertise. Furthermore, most pathologists follow simple algorithmic decision tree approaches that use only limited amounts of information presented in these images [6]. For the same task on lung cancer, Yu et. al [7] used an automatic image-segmentation pipeline to identify the tumor nuclei and tumor cytoplasm from which they extracted image features. Several classical machine learning approaches have been developed to achieve classification of lung [8,9] and breast [10,11] cancers. Advances in deep learning have paved the way for artificial neural networks to surpass most traditional machine learning methods in the field of image processing. In particular, convolutional neural networks (CNNs) consists of convolutional layers which exploit locality and stationarity and were first proposed by Le Cun et al. [12] These approaches have quickly risen to the state-of the-art on almost all image-based tasks including medical imaging [13, 14]. One of the benefits of using a convolutional neural network architecture is that using domain knowledge to handcraft an image feature extraction system is not required. Conveniently, feature extraction is an automatic task for neural models, and is trained end-to-end in a CNN [15]. In recent years, a large number of high resolution digital Whole Slide Images (WSI) of H&E stained tissue slides have been captured by pathologists [16]. With an increasing amount of “big data”, CNN has become an excellent candidate for WSI processing tasks and it has been adapted by many studies already for tasks like prediction of kidney function and segmentation [17,18], breast cancer detection [19], [43], tumor prediction for osteosarcoma [20], colon cancer detection [21] and analysis of tumor-infiltrating lymphocytes [42]. In 2014 the GoogLeNet [22] architecture won the Imagenet Large Scale Visual Recognition Challenge Since then, it has been adapted for a wide range of medical imaging studies [23–25], such as for breast and skin cancer detection where it outperforms human experts. Khosravi et al. [26] compared a basic CNN architecture with different versions of Inception and Resnet architectures for classifying pathology images. They obtained an accuracy of %100 and %79 for intra and inter images for tumor subtype discrimination using fine-tuned Inception-V3 on TCGA lung data [27]. Coudray et al. [28] trained an implementation of Inception V3 on the lung cancer classification task (inter-images) and obtained a macro AUC of 0.97 (AUC ∼0.993 for distinguishing between neoplastic and non-neoplastic tissue, and 0.947 for distinguishing between the neoplastic subtypes). The high performance of these Convolutional Networks architectures comes with a high computational cost as they require extremely large numbers of parameters and operations for this level of accuracy which makes the deployment challenging [29,30]. Therefore, software optimization and efficient architecture design becomes a key for many deep learning applications to address the trade-offs between performance and efficiency. In this work, we implement, train and evaluate a simple CNN architecture named PathCNN. We show that the proposed architecture converges faster, uses less memory and achieves better performance than complex architectures. Importantly, PathCNN’s more efficient use of computational resources, allowed us to simultaneously train on images from multiple tumor sites, subtypes and corresponding normals. This in turn, uncovered several cases of misclassified images, outliers (e.g. fibrosis, tubular atrophy, calcification) as well as staining irregularities and other artifacts in the TCGA dataset. In short, PathCNN can be applied to quickly classify whole-slide images and detect outliers in the data.

## Materials and Methods

### Dataset compilation

High-resolution (20x magnification, 0.5um per pixel) tissue whole-slide images of lung cancer, kidney cancer, breast cancer and non-neoplastic (normal) tissue were downloaded from the Genomic Data Commons Database from The Cancer Genome Atlas (TCGA) [31, 32]. The following datasets were used for this study: the lung cancer dataset was composed of: 680 slides of Lung Squamous Cell Carcinoma (TCGA-LUSC), 733 slides of Lung Adenocarcinoma (TCGA-LUAD) and 494 Normal tissue slides; the kidney cancer dataset was composed of 147 images of Kidney Chromophobe Renal Cell Carcinoma (TCGA-KICH), 896 images of Kidney Clear cell Renal Cell Carcinoma (TCGA-KIRC), 345 images of Kidney Papillary Renal Cell Carcinoma (TCGA-KIRP) and 659 Normal tissue images; the breast cancer dataset was composed of 1412 images of Breast Invasive Carcinoma (TCGA-BRCA) and 337 Normal tissue images. Our overall computational strategy is summarized in **Figure 1** and is further explained in the following sections.

**Figure 1.**
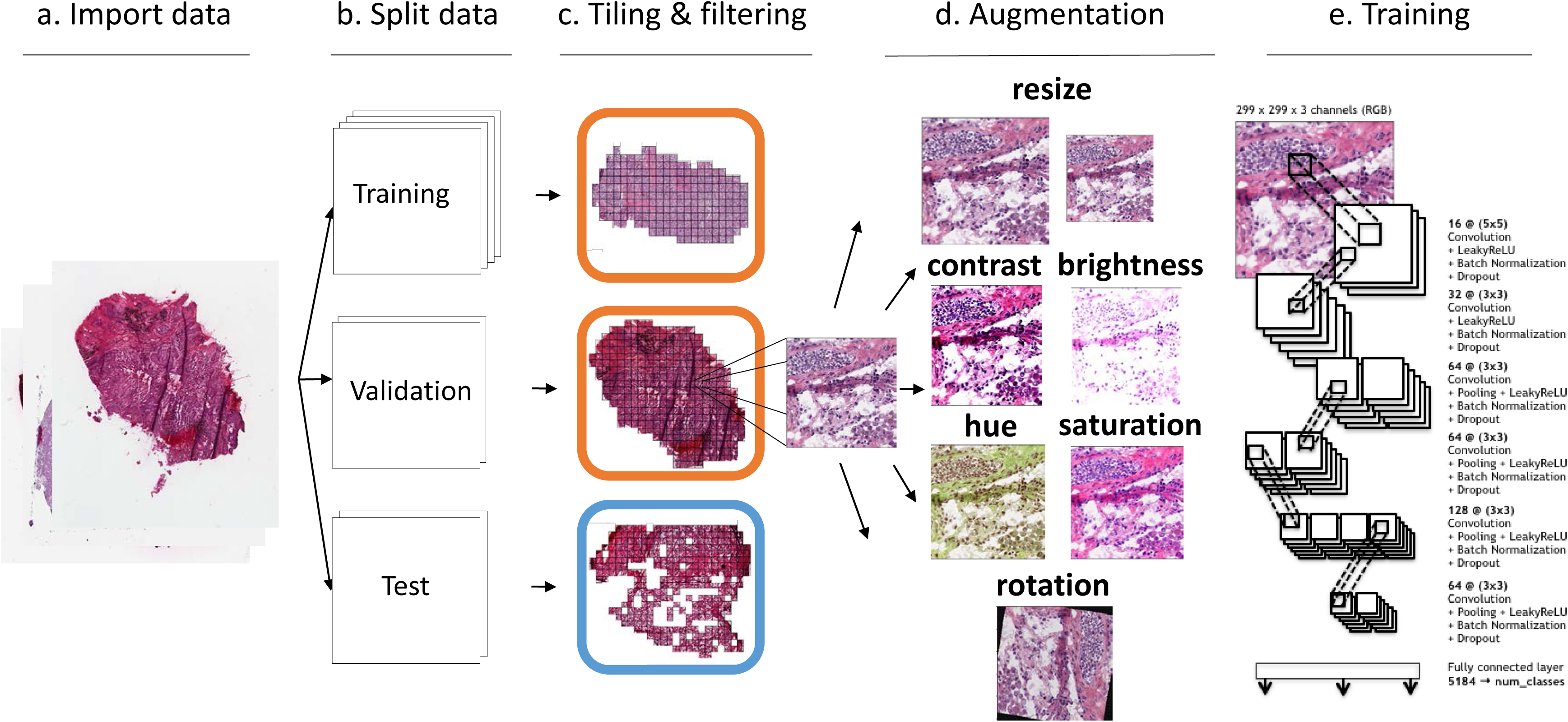
Data and Workflow. **(a)** Whole-slide images of resected patient tumors were downloaded from the Genomic Data Common database, **(b)** Slides were then split into a training (70%), a validation (15%) and a test set (15%); slides from the same patient were included in only one of the three sets, **(c)** Slides were tiled by non-overlapping 512×512 pixels windows, omitting those with over 50% background, **(d)** Training data was randomly augmented using different approaches, **(e)** Architecture of the Convolutional Neural Network used for training.

### Data pre-processing

The whole-slide images were tiled into non-overlapping 512×512 pixel tiles using open source package openslide [33]. This tiling strategy is commonly employed for high-resolution medical images, otherwise the resolution would be limited by the GPU memory [34]. 753 slides (94 lung, 430 kidney and 229 breast) initially uploaded were removed because of compatibility and readability issues. Additionally, tiles with more than 25% of background were filtered out, as they are unlikely to contain informative features. This process generated nearly 665,812 tiles for lung, 1,077,076 for kidney and 655,885 for breast. Before feeding the images into the network, we further down-sampled the tiles to 299 × 299 pixels to make the network comparable to Google’s Inception V3, which requires 299 × 299 pixel inputs.

### Data split into training, validation and test sets

For the three cancer types, the data was split into 70%, 15% and 15% into training, validation and test datasets respectively (**Table 1**). To avoid leakage, we ensured that all slides (and tiles) that came from the same patient were included in the same dataset (training, validation, or test). This ensured that the model could not simply learn histologic features that are patient-specific to achieve good performance at validation and test time.

**Table 1:** Number of tiles and slides in the lung, kidney and breast datasets.

|  | **Training**<br>**(tiles/slides)** | **Validation**<br>**(tiles/slides)** | **Testing**<br>**(tiles/slides)** |
| --- | --- | --- | --- |
| LUAD | 176,839/513 | 42,485/111 | 40,063/110 |
| LUSC | 218,260/473 | 46,687/104 | 41,969/103 |
| Normal lung | 339,572/460 | 81,303/99 | 73,979/100 |
| KICH | 98,817/102 | 21,665/23 | 20,059/22 |
| KIRC | 221,023/621 | 60,147/139 | 57,372/136 |
| KIRP | 68,114/242 | 20,162/49 | 14,863/54 |
| Normal kidney | 66,724/349 | 16,700/82 | 16,085/83 |
| BRCA | 421,467/988 | 96,992/211 | 92,461/213 |
| Normal breast | 26,091/211 | 10,064/63 | 8,810/63 |

### Data augmentation

Data augmentation creates new sample points by making alterations to the existing dataset and is widely used to increase the robustness of the image classifier [8,35,36]. It is a form of regularization, as it helps highly complex neural networks to avoid over-fitting by creating more training examples for the network to learn from. The rotational orientation does not change the classification of the tissue and varying dying methods and lighting environments means that the same histological identifying features may be slightly different in color and saturation between slides. Therefore, using the Tranforms library from Torchvision [37], along with self-written functions and classes, we augmented the training data by randomly introducing positional transforms such as: resize, a horizontal flip, or a rotation of 0°, 90°, 180° or 270° degrees. Additionally, we randomly adjusted the hue, brightness, contrast, saturation of the image as visualized in **Figure 1d** and normalized ((image - mean) / std) before passing them to the model.

### Efficient deep learning architecture development

A typical ResNet type block of layers for convolutional neural networks is: Convolution → Non-linearity → Batch Normalization → Pooling → Dropout [26]. Our network uses the same construction, that is similar to the first few layers of the Google Inception V3 model. We initially used 3 convolution blocks going from 32 → 64 → 80 features. These blocks did not include a pooling function, so we added a final linear layer to map 25,920 features down to the number of classes. However, we found that without pooling the features together throughout the blocks, the model was unable to properly perform inference, which may be why the Inception model follows these upsampling blocks with several layers of pooling blocks. Therefore, we tried a 4 convolutional block network going from 16 → 32 → 64 → 32 output features, using pooling in every block. With this simpler architecture, the model was able to learn, but we were only able to obtain a macro AUC of ∼0.903 on the validation set. We concluded that a deeper network may better represent the complexity of the problem. The final architecture we developed can be visualized in the **Figure 1**. We used 6 layers of convolutional blocks 16 → 32 → 64 → 64 → 128 → 64, followed by a fully connected layer that mapped 5184 features down to the number of classes. The first block uses a kernel size of 5, while the following blocks all used a kernel size of 3. Furthermore, the first two blocks did not contain a pooling layer, as we wanted to upsample the number of features before downsampling with max pooling.

### Hyper-parameter tuning

In our experiments, we found that tuning the dropout rate was very important for model performance. With a dropout rate of p = 0.5, we were only able to obtain an AUC of ∼0.857 on the validation set, while when we decreased dropout to 0.1, we were able to consistently reach macro AUC above 0.9. For aggregation, we used the average score across all the tiles for each class which outperformed the proportion method. For nonlinearity, we used leaky relu as it performed slightly better than relu only. For hyper-parameter tuning we performed a search similar to coordinate descent algorithm, where we varied a particular hyper-parameter until we found the best performing model, then moved on to optimize the next hyper-parameter. After several iterations to find a proper learning rate, we found that the best learning rate was 0.001. As for the initialization, Xavier initialization helped get the model achieve better performance earlier (AUC 0.94 after 1 epoch vs 0.89) compared to Gaussian initialization. For data augmentation, we used random horizontal flip, random rotations and random change in brightness, contrast, saturation and hue of an image. We found significant boost in performance from AUC of 0.93 to above 0.96. As for the optimizers, both Adam and RMSProp outperformed SGD; we used Adam in our final architecture. For more details, see Supplemental Information.

### Inception V3 architecture

To assess the performance of our architecture, we replicated the work done by Coudray et al. [28] on lung cancer task using the pipeline publicly available at [39]. To this end, we used the same training loop and aggregation methods, but with Google’s Inception V3 architecture. Inception increases representational power for image classification by using” Network in Network” approach [38] and reduces computational needs [8]. The architecture consists of blocks of a Stem (Convolution → Convolution → Convolution padded → MaxPooling → Convolution → Convolution → Convolution) → 3x Inception → 5 x Inception → 2 x Inception → AvgPooling → Dropout(40%) → Fully connected → Softmax with blocks going from 32 → 32 → 64 → 64 → 80 → 192 → 288 → 768 → 1280 → 2048 → 1000 features. [22]. The inception layer combines convolution blocks with various filter sizes and pooling layer which helps with classification when there is a large variation in the location and the size of the information presented in the image [44].

### Tile aggregation

While the neural network was trained on a per-tile basis, the model must perform classification on a per-slide basis. Thus, the per-tile scores from the neural network were aggregated for the final prediction using two different methods: (1) per-slide average score, where the average score across all the tiles for each class is calculated, and (2) per-slide tile proportion, where the class with the maximum score is assigned to each tile. The final score is the proportion of tiles assigned to each class.

### Model evaluation metrics

The Area Under the Curve (AUC) was used as the evaluation metric. AUC is traditionally defined only for binary classification, but the lung and kidney tasks had more than two classes. Therefore, first the AUC of individual classes using a one-vs-all method were calculated. Then, the macro-average (macro-AUC), which is the unweighted mean of the individual class AUCs were calculated. Additionally we calculated the following metrics to further evaluate the performance and the robustness of our results: Average of Precision and Recall; Cohen’s kappa (Eq.1), an indicator of agreement of results commonly used for multi-class and imbalanced class problems; Jaccard Coefficient (Eq.2), a measure of similarity and calculated by Intersection over Union of predicted and true labels; Log-loss/cross entropy, the negative log-likelihood of the true labels given a probabilistic classifier’s predictions [27] and it indicates much the predictions vary from the actual labels.

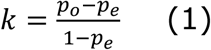

Where p_o_is relative observed agreement and p_e_is the hypothetical probability of chance agreement between the predictions and the actual labels.

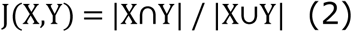

Where X and Y are the prediction and actual label set.

### Pan-cancer t-SNE visualization

For the pan-cancer analysis, we used our models’ representation of each image in later layers before reducing them into class probabilities, then clustered those representations using Distributed Stochastic Neighbor Embedding (t-SNE). To achieve this, we used the last layer (fully connected layer) weights which embeds the information in an array of size 5184 that is later reduced to class probabilities to predict the label. We only extracted the weight that belongs to tiles that were correctly classified by the model to represent the given image, to increase the confidence interval of the clustering. Then t-SNE was used to visualize these high dimensional embedding (number of slides x 5184) in 2-dimensional space.

### Data and code availability

Code and Jupyter notebooks to help reproduce the analysis are available at https://github.com/sedab/PathCNN.

## Results

### Simplified deep learning model achieves high accuracy and fast convergence

To evaluate our deep learning architecture, we assessed its performance on the lung dataset (LUAD, LUSC and normal lung from TCGA) and directly compared to the state-of-the-art Inception V3 architecture previously used for this task [37,28]. When there was no time constrains, we were able to obtain a macro AUC of 0.979 for the TCGA validation set at epoch 10 (when the model fully converged) and an AUC of 0.957 for the TCGA test set on our baseline lung cancer task, comparable to previously reported AUC achieved by Inception V3 [28]. Detailed comparison of macro AUCs between Inception and PathCNN as a function of the number of steps on validation set can be seen in **Figure 2a**. The model does a nearly perfect job when the classification task is only between tumor and normal lung tissue, with AUC of 0.99. Differentiating between the two types of lung cancer, LUAD and LUSC, does not perform equally well, with individual one-vs-all AUC of 0.93 for both classes (**Table 2**). This agrees with previously published results [27, 28] showing that tumors are visually distinct from normal tissue, while cancer types or subtypes can be more challenging to visually differentiate. The results from Yu et al. [7] also follow a similar pattern with a significant difference in AUC between the two tasks. Examples of predictions of our model and Inception V3 on selected lung adenocarcinoma (LUAD) and lung squamous cell carcinoma (LUSC) slides are visualized using heat-maps (**Supplementary Figure 1**).

**Figure 2.**
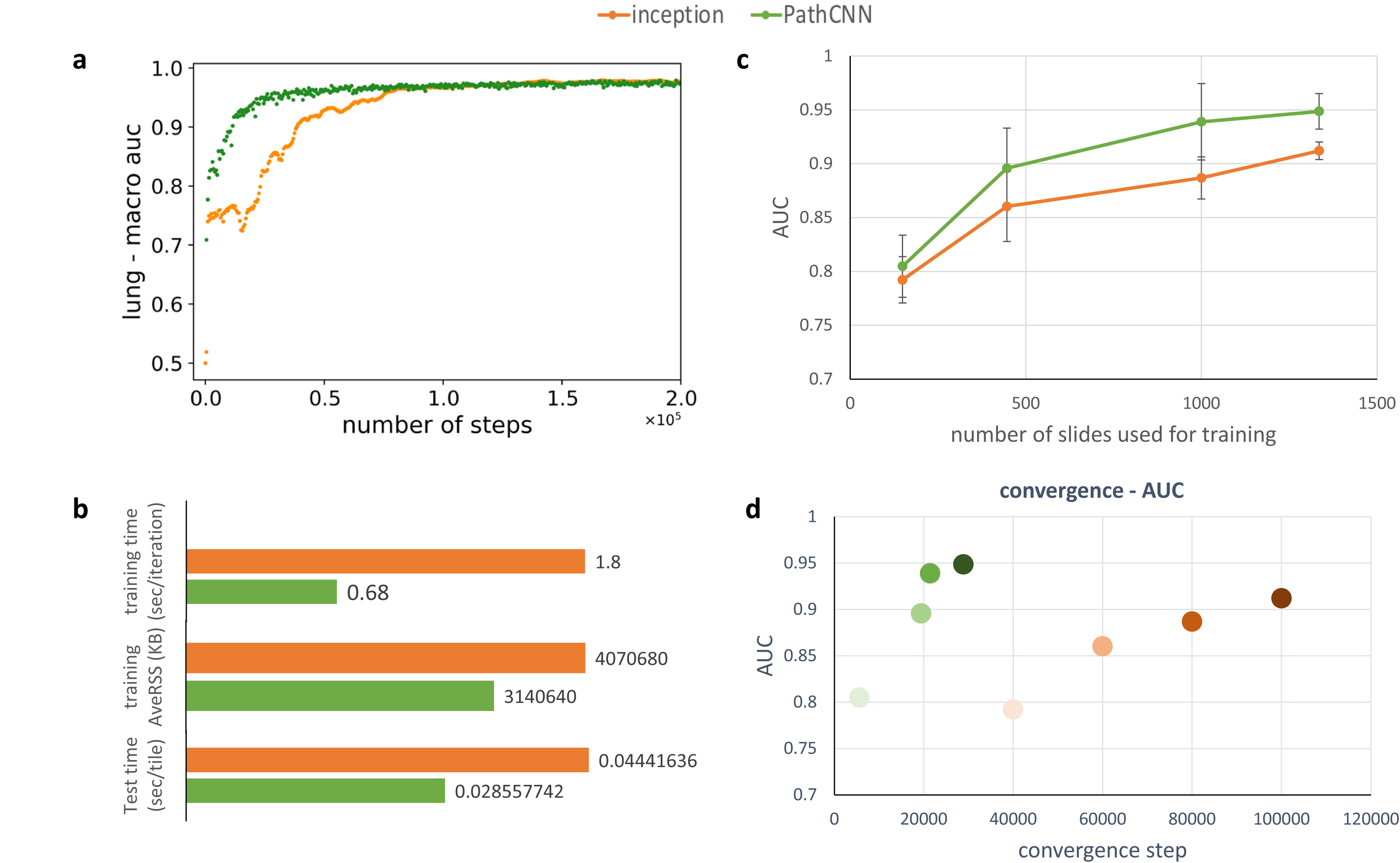
Evaluation of PathCNN and comparison with Inception v3. **(a)** Macro AUC at each training step on TCGA training dataset, **(b)** Computing resource requirements: training time (seconds per iteration), training memory (Gb) and test time (seconds per tile), **(c)** Impact of the size of the training dataset on the test AUC**, (d)** Convergence time (x-axis) vs AUC (y-axis) for each architecture and training set size (darker color represents larger training set). Color assignments: Green=PathCNN and Orange=Inception.

**Table 2:**
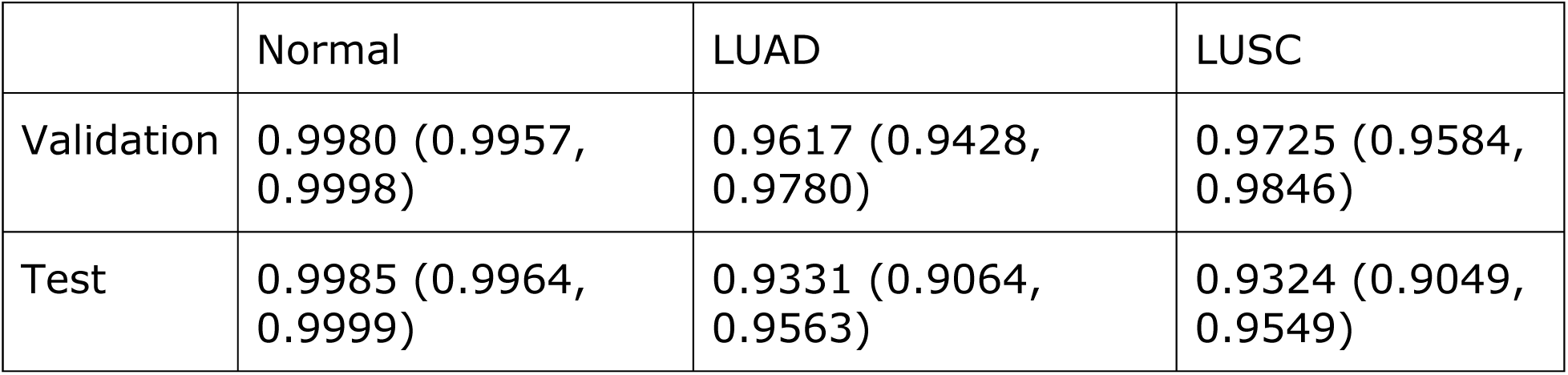
Performance of lung type classification models. Validation and Test Macro AUCs (with 95% confidence interval) for the proposed model and for Inception V3.

### Memory usage and speed comparison

Both models were trained on NVIDIA P100 GPU with 100GB memory. During training, our model was much faster compared to inception (**Figure 2b**): our model required only 0.68 sec per iteration, whereas inception needed 1.8 sec per iteration. Furthermore, the average memory occupied at RAM was lower than inception (3GB vs 4GB). Also, our model is much faster at test time: 0.29 sec per tile compared to 0.044 sec with inception, i.e. more than 6 times faster.

### Evaluation on independent cohorts

We then tested the performance of both models on independent datasets collected the NYU Medical Center [28]. More specifically, we tested the models on two types of specimens: surgical resections (98 frozen slides and 140 FFPE slides) and biopsies (102 slides). ROC curves of the performance of our model on each of the three collections of whole-slide images are drawn in **Supplemental Figure 2**. The performances are overall comparable to those published previously: the LUAD/LUSC AUC on frozen slides is slightly lower (0.85/0.90 for PathCNN vs 0.91/0.94 for Inception), while the ones on FFPE (0.91/0.97 vs 0.83/0.93) and Biopsies (0.86/0.90 vs 0.83/0.86) are greater.

### Impact of training dataset size on performance

To measure how the amount of data used effect the model performances, we performed a down-sampling analysis as follows: we trained the model on reduced datasets by sampling slides (sampled at 1/9th, 1/3rd and 3/4^th^ after ordering by name) and the full set (number of slides = 149, 446, 1000 and 1335) and tested on the same full test set from TCGA. Then we calculated the average of AUCs obtained by testing on checkpoints after down-sampled datasets validation sets performance converged during training. As summarized in **Figure 2c**, we found that PathCNN generalized to the external dataset much better for when trained on smaller datasets compared to Inception (p = 0.0952, 0.0027, 0.0025, 2.6E-14). Furthermore, our model had much faster convergence and showed better convergence-AUC trade-off for the external datasets (**Figure 2d**).

### Impact of augmentation on performance

We found that the augmentation had a big effect on model performance. AUC increased by ∼0.02 (p<0.001) on the TCGA test data set (**Supplemental Figure 3**). Furthermore, interestingly we observed faster overfitting when the training data was not augmented.

### Performance on kidney and breast whole-slide images

We then retrained our model on all TCGA datasets: lung, kidney and breast. Although optimized on the lung cancer dataset, our model transferred quite well to the kidney and breast classification tasks. While we did retrain the entire network (in contrast to a transfer learning approach), we found that even without re-tuning the hyper-parameters, the AUCs were very high (ROC curves are shown in **Figure 3a-c**). In fact, we did not see the same kind of drop in performance between the validation and tests sets, and they were able to achieve a macro-AUC around 0.998 for kidney (in 12 epochs) and 0.994 for breast (in 14 epochs) (**Table 3** and **Table 4**).

**Figure 3.**
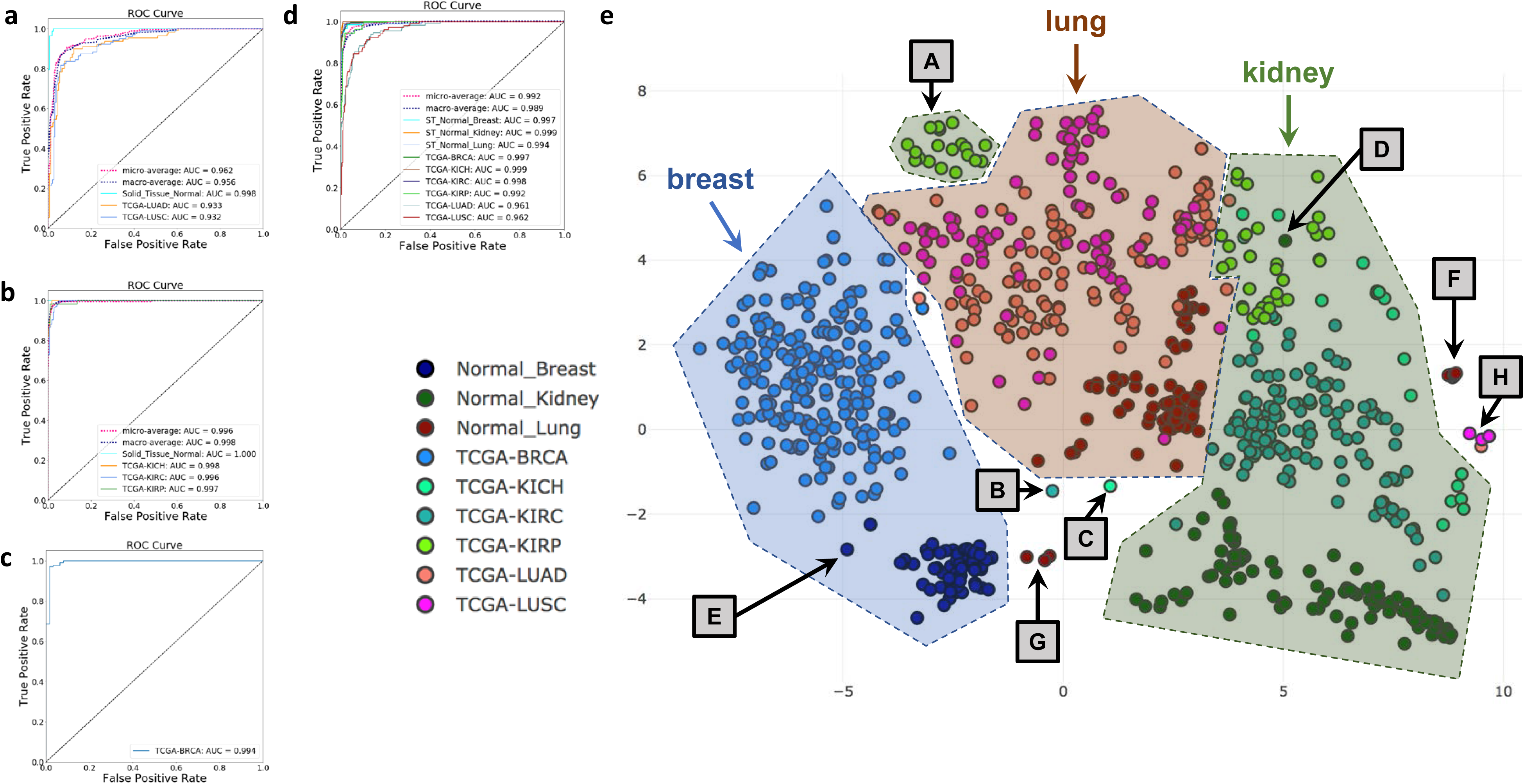
PathCNN detects misclassified slides and staining artifacts in TCGA. **(a)** ROC curve on the test set of lung images (normal lung, LUAD and LUSC), **(b)** ROC curve on the test set of kidney images (normal kidney KICH, KIRC and KIRP), **(c)** ROC curve on the test set of breast images (normal breast and BRCA), **(d)** ROC curve for all slides in the test set, **(e)** t- SNE visualization of the PathCNN model trained on all cancer types and corresponding normal (lung, kidney and breast); detailed description of the identified clusters in Table 7.

**Table 3:** Validation and test performance of classification into lung, kidney and breast. Validation and Test Macro AUCs (with 95% confidence interval) for the proposed model.

|  | Lung | Kidney | Breast |
| --- | --- | --- | --- |
| Validation | 0.9784 (0.9503, 0.9993) | 0.9986 (0.9953, 1.0000) | 0.9992 (0.9989, 0.9995) |
| Test | 0.9565 (0.9093, 0.9997) | 0.9980 (0.9929, 1.000) | 0.9937 (0.9935, 0.9980) |

**Table 4:** Validation and test performance of classification into kidney classes. Validation and Test Macro AUCs (with 95% confidence interval) for the proposed model.

|  | Normal | KICH | KIRC | KIRP |
| --- | --- | --- | --- | --- |
| Validation | 0.9999 (0.9998, 1.000) | 0.9983 (0.9960, 1.0000) | 0.9973 (0.9942, 0.9996) | 0.9980 (0.9954, 0.9998) |
| Test | 0.9990<br>(0.9976,<br>0.9999) | 0.9935<br>(0.9867,<br>0.9989) | 0.9940<br>(0.9891,<br>0.9975) | 0.9917<br>(0.9837,<br>0.9974) |

Lastly, the performance of our algorithm is evaluated using various statistical metrics: AUC, average of Precision and Recall, Cohen’s kappa, Jaccard Coefficient, and Log-loss. We found high degree of agreement and similarity between the model class predictions and the actual classes as indicated by the Cohen’s kappa and Jaccard coefficient in **Table 5**. Similar to precision, recall and log-loss we see higher values with breast followed by kidney and lung.

**Table 5:**
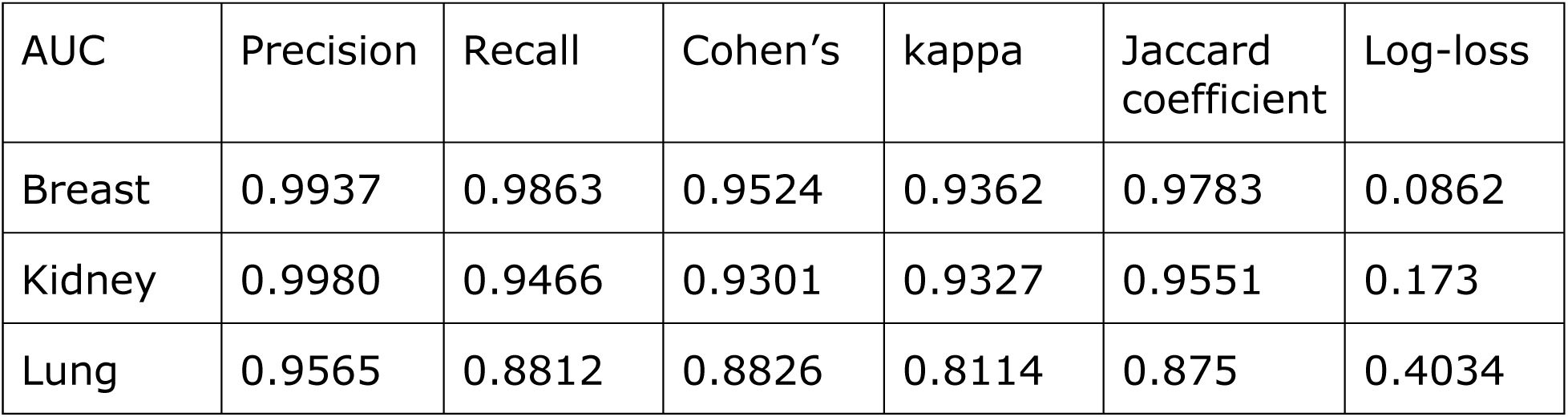
Performance evaluation using various statistics measures.

| AUC | Precision | Recall | Cohen’s | kappa | Jaccard coefficient | Log-loss |
| --- | --- | --- | --- | --- | --- | --- |
| Breast | 0.9937 | 0.9863 | 0.9524 | 0.9362 | 0.9783 | 0.0862 |
| Kidney | 0.9980 | 0.9466 | 0.9301 | 0.9327 | 0.9551 | 0.173 |
| Lung | 0.9565 | 0.8812 | 0.8826 | 0.8114 | 0.875 | 0.4034 |

### t-SNE visualization of the breast-kidney-lung model and outlier detection

We then used PathCNN to simultaneously train for all the breast, kidney and lung types/subtypes and corresponding normals (BRCA, KIRP, KIRC, KICH, LUSC, LUAD and normal breast, kidney, lung). We obtained macro AUC of 0.987 on the validation set and 0.989 on the test set. ROC curves can be seen in **Figure 3a-d**. We then performed dimensionality reduction on the weights of the last fully-connected layer using t-Distributed Stochastic Neighbor Embedding (t-SNE) [45] and visualized the result in two dimensions (**Figure 3e**, each dot represents a whole-slide image). Overall, we observed a clear separation by organ (breast, kidney or lung), and by cancer type or normal within each organ (see blue, green and brown areas in **Figure 3e**). However, we also noticed smaller clusters of slides (clusters A through H) outside these three large areas. We then asked a pathologist to examine all these slides and annotate them. These slides and the pathologist’s annotations – grouped by slide type – are listed in **Table 6**. More specifically, area A represents a cluster of KIRP cases characterized by intense basophilic staining. Area B is a single KIRC outlier (with a slide composed of predominantly fat) and area C a KIRP outlier (that shows significant frozen artifact). Area D is a normal kidney slide with interstitial fibrosis, tubular atrophy and calcification-reactive features that can also be seen among KIRP cases. Area E includes two normal breast outliers: these slides were found to be overstained.. In area F, the pathologist found normal lung outliers, mainly hypereosinophilic, while one case represented a sample from a benign collapsed lung. Additional normal lung outliers were found in area G, with intense basophilic staining, and one case of benign lung tissue with emphysematous change. Lastly, area H comprises LUAD and LUSC outliers, mainly characterized by a washed out staining pattern. Representative examples of slides belonging to each cluster are shown in **Supplemental Figure 4**.

**Table 6:** Annotation of t-SNE outliers by pathologist.

| Area | Discrepancy | Sample ID | Interpretation |
| --- | --- | --- | --- |
| A | KIRP cluster | TCGA-B1-A655-01A-01-TS1 | hyperbasophilic staining |
|  |  | TCGA-BQ-5885-01A-01-BS1 | hyperbasophilic staining |
|  |  | TCGA-2Z-A9JQ-01A-01-TS1 | hyperbasophilic staining |
|  |  | TCGA-2Z-A9JE-01A-01-TS1 | hyperbasophilic staining |
|  |  | TCGA-UZ-A9PV-01A-01-TSA | hyperbasophilic staining |
| B | KIRC outliers | TCGA-B8-A8YJ-01A-01-TS1 | predominantly fat |
| C | KIRP outlier | TCGA-KL-8337-01A-01-BS1 | frozen artifact |
| D | Normal kidney among KIRP | TCGA-GL-8500-11A-01-TS1 | significant interstitial fibrosis, tubular atrophy, calcification |
| E | Normal breast outliers | TCGA-BH-A0C3-11A-01-TSA | overstained slide |
|  |  | TCGA-BH-A1ET-11B-02-TSB | overstained slide |
| F | Normal lung outliers | TCGA-22-4593-11A-01-TS1 | hypereosinophilic staining |
|  |  | TCGA-73-4659-11A-01-BS1 | hypereosinophilic staining |
|  |  | TCGA-73-4670-11A-01-BS1 | hypereosinophilic staining |
|  |  | TCGA-49-4510-11A-01-TS1 | benign collapsed lung |
|  |  | TCGA-22-5477-11A-01-TS1 | hypereosinophilic staining |
| G | Normal lung outliers | TCGA-44-6775-11A-03-TS3 | emphysematous change |
|  |  | TCGA-44-5645-11A-03-TS3 | hyperbasophilic staining |
|  |  | TCGA-44-5645-11A-02-TS2 | hyperbasophilic staining |
| H | LUAD outlier | TCGA-86-7714-01A-01-TS1 | washed out |
| H | LUSC outliers | TCGA-85-8664-01A-01-TS1 | washed out |
|  |  | TCGA-85-8666-01A-01-TS1 | washed out |
|  |  | TCGA-85-8288-01A-01-TS1 | washed out, large pieces of cartilage |

## Discussion

With the increasing number of accumulated WSIs, there is a significant opportunity for developing effective diagnostic tools using Machine Learning techniques to exploit visual features in pathology images that are not detectable by the human eye [40]. Recent developments in computer vision has opened new doors for the development of such tools which comes with the expense of computational requirements. Using our simple convolutional neural network approach, we have exceeded the performance of Inception V3 network [7] trained for one week on the lung cancer classification task with a macro AUC of 0.957. This difference is reflected in the reduced training time of 2 days as compared to 1 week for the deep network. This also means it will be significantly faster at prediction time; aggregation and prediction for a particular slide image took around 30 minutes. Limited resources in a realistic health care setting means that this decrease in diagnostic turnaround time could translate to positive real-world outcomes for patients. We found that using data augmentation, tuning the learning rate, reducing the drop-out rate, and increasing the depth were the most important factors for achieving the best performance. With a simpler architecture, we were able to reach comparable results to a very deep network. With further hyper-parameter tuning and further software optimization, better results and higher efficiency can be achieved. For lung cancer, the individual class AUCs reflect how the network was better at distinguishing between Solid Tissue Normal (non-neoplastic tissue) and the two types of non-small cell lung cancer, with an AUC of around 0.999. On the other hand, the network’s performance is not as good when trying to differentiate between the two different classes of cancer: Lung Squamous Cell Carcinoma (TCGA-LUSC) and Lung Adenocarcinoma (TCGA-LUAD). Another positive outcome from our work is that it generalizes well to kidney and breast cancers. In fact, the AUC scores improved in comparison to the ones achieved for lung: macro AUC of 0.997 and 0.996 on the test set respectively. As with lung cancer, the model performed significantly better at distinguishing non-neoplastic tissue from neoplastic tissue for both of these cancer types. For breast cancer, there were only two possible classes: non-neoplastic and neoplastic tissue. Just as the case with lung cancer, we found that our model can easily distinguish between these two classes even though we did not further tune the hyper parameters specific to this data set. This suggests that the framework we developed is well suited for the general task of distinguishing neoplastic tissue from non-neoplastic tissue. For the kidney cancer task, there were more neoplastic subtypes (3 compared to 2 for lung). Nevertheless, we found that the model was almost perfectly able to distinguish between them and achieved a macro AUC better than the lung cancer task. On the test set, the individual class AUCs were 0.999, 0.994, 0.994, and 0.992 for Solid Tissue Normal, TCGA-KICH, TCGA-KIRC, TCGA-KIRP, respectively. This is likely due to the fact that these neoplastic subtypes are more visually distinct than the lung cancer subtypes, and therefore the task is easier overall. Another possible explanation could be due to the difference in the available training data. After tiling, we had around 1077k (925 slides) kidney cancer tiles compared to 666k (1928) lung cancer tiles in their respective training sets, a 62% increase. This phenomenon may be explained by the pre-processing step that removed any tile with more than 25% background. The kidney tissue slides may be denser with richer information.

## Acknowledgments

We would like to thank the Applied Bioinformatics Laboratories (ABL) for providing bioinformatics support and helping with the analysis and interpretation of the data. ABL is a shared resource, partially supported by the Cancer Center Support Grant, P30CA016087, at the NYU School of Medicine Laura and Isaac Perlmutter Cancer Center. This work has used computing resources, services and staff expertise at both the NYU Medical Center and NYU IT High-Performance Computing Facilities. The slide images and the corresponding cancer information were uploaded from the Genomic Data Commons portal https://gdc-portal.nci.nih.gov and are in whole or part based upon data generated by the TCGA Research Network http://cancergenome.nih.gov/). The data used were publicly available without restriction, authentication or authorization necessary.

---

**Supplemental Figure 1.**
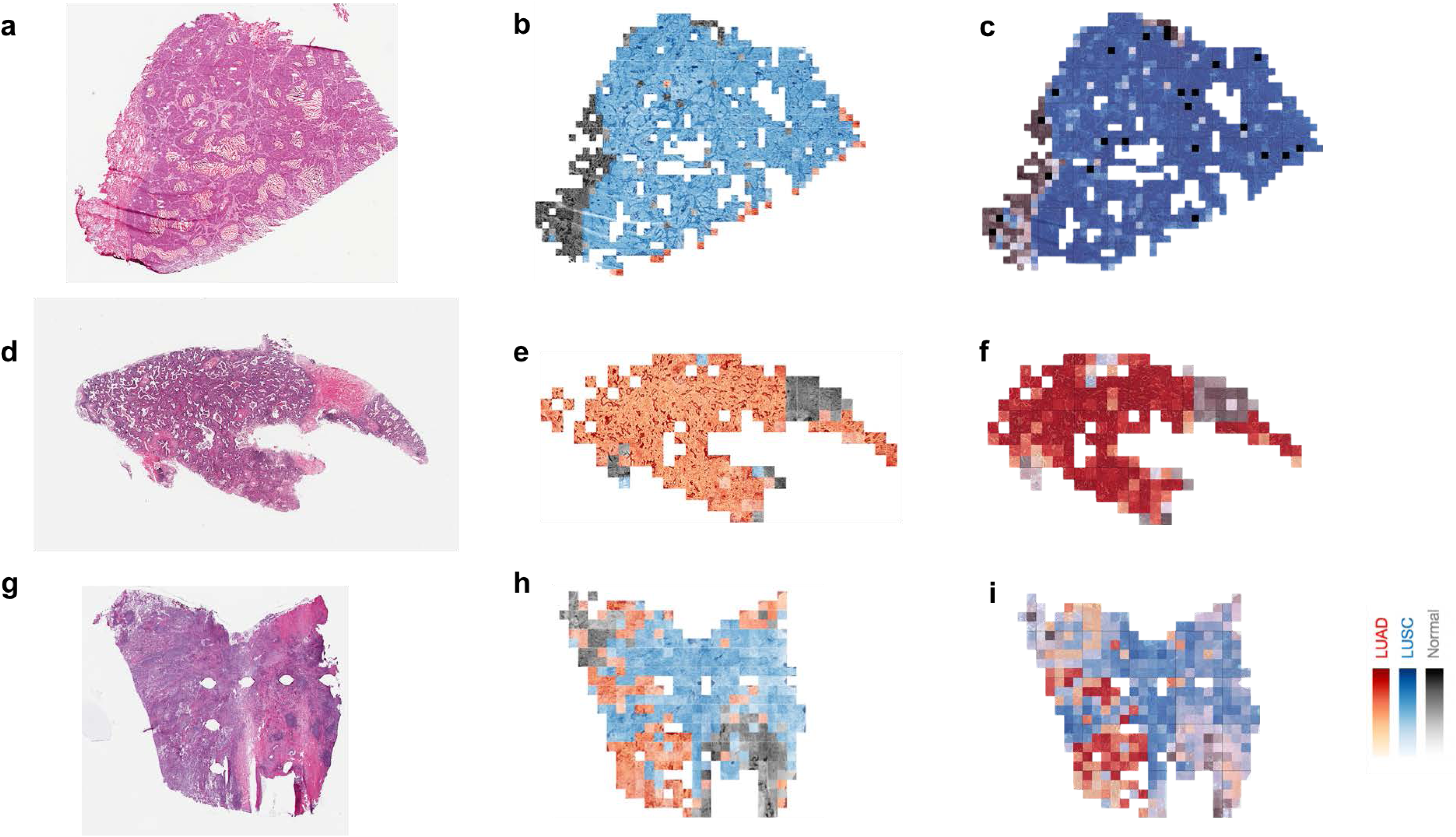
Visualization of images and classification probability heatmaps. **(a)** Original Whole Slide image with Lung Squamous Cell Carcinoma (TCGA-LUSC), **(b)** Aggregated prediction using tiles of the slide for the proposed model, **(c)** Aggregated prediction using tiles of the slide for Inception V3, **(d)** Original Whole Slide image with Lung Adenocarcinoma (TCGA-LUAD), **(e)** Aggregated prediction using tiles of the slide for the proposed model, **(f)** Aggregated prediction using tiles of the slide for Inception V3, **(g)** Original Whole Slide image with Lung Adenocarcinoma (TCGA-LUAD), **(h)** Aggregated prediction using tiles of the slide for the proposed model, **(i)** Aggregated prediction using tiles of the slide for Inception V3.

**Supplemental Figure 2.**
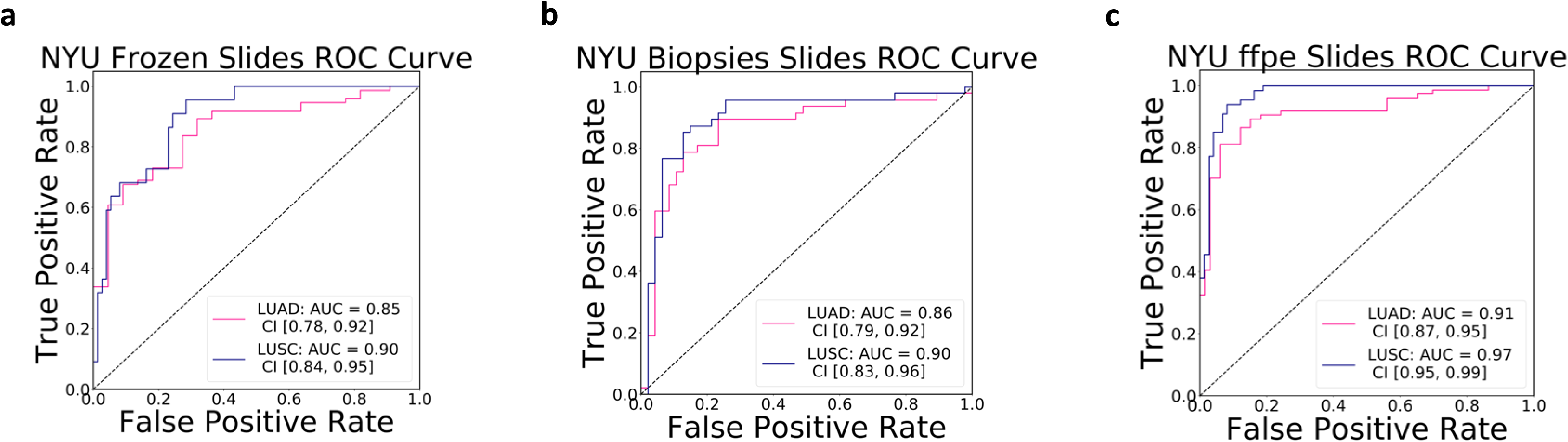
PathCNN performance on independent NYU cohort. **(a)** ROC curve on slides prepared from tumor resections (Frozen), **(b)** ROC curve on slides prepared from biopsies (Frozen and FFPE), **(a)** ROC curve on slides prepared from tumor resections (FFPE).

**Supplemental Figure 3.**
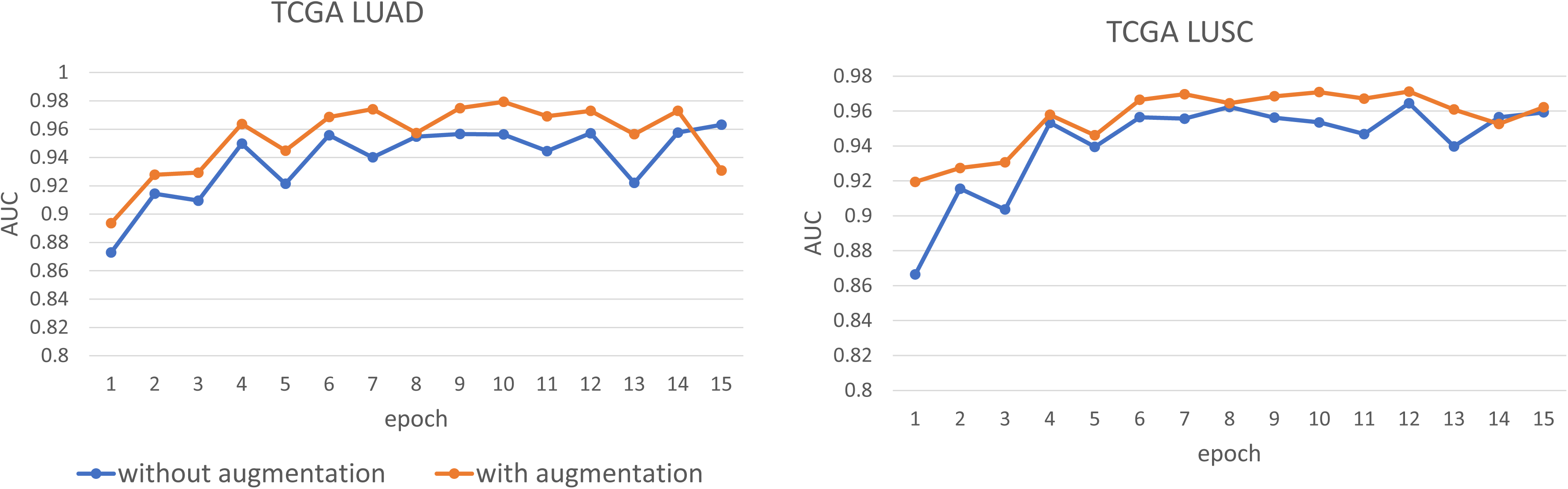
Impact of augmentation on PathCNN performance. Macro AUC at each training step on TCGA training dataset. Color assignments: Blue=no augmentation, Orange=augmentation enabled.

**Supplemental Figure 4.**
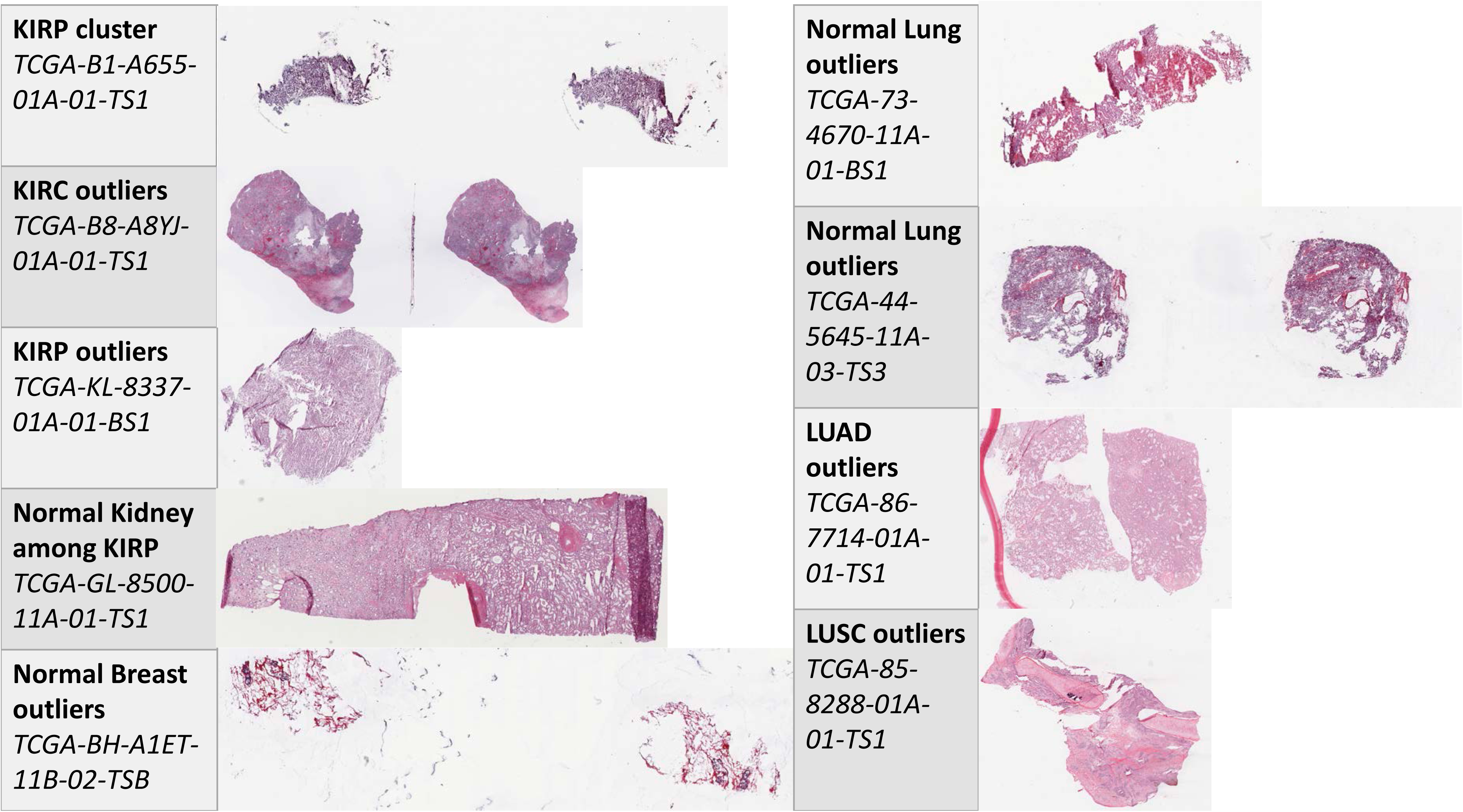
Representative slides from outlier analysis. Representative slides from areas A through H marked on the t-SNE plot of Figure 3e.

## Supplemental Information

### Nonlinearity

In our experiments, we found that both ReLU and LeakyReLU worked well, but using LeakyReLU resulted in slightly better performance. Therefore for our final architecture, we used LeakyReLU with negative slope 0.01 as the non-linearity.

### Dropout

In our experiments, we found that tuning the dropout rate was very important for model performance. When we first used the default dropout rate of p = 0.5, we were only able to obtain an AUC of $\sim$0.857. We realized that the dropout rate may be too high, especially as we were applying dropout after every convolutional block. In comparison, the Inception model only applies one layer of dropout after all of the blocks. When we decreased dropout to 0.1, we were able to consistently reach macro AUC above 0.9. Another explanation may also be due to the use of batch normalization within each convolutional block. Both batch normalization and dropout are regularization methods, so the use of batch normalization reduces the need for dropout. In this case, the higher dropout of $p = 0.5$ was causing our model to underfit.

### Aggregation method

We tried both the average score and proportion of tiles methods for aggregating the tiles, as described in the methodology section 3.7.1. We found that the methods were comparable, though the average method performed between 0.003 and 0.008 AUC better on the three datasets. Intuitively, this makes sense: Assigning each tile a class based on the maximum probability would treat a tile that is assigned a class with 99\% probability the same as one with 60\% probability. Therefore, taking a “majority vote” method per tile could potentially discard some useful information for the prediction.

### Other hyperparameters

We didn’t run a full grid search on all the hyperparameters and options listed in the methodology section. However, we performed a search similar to coordinate descent algorithm where we varied a particular hyperparameter until we found the best performing model, then moved on to optimize the next hyperparameter. With this method, we were still able to draw some insights into the effect of particular hyperparameters on model performance. One of the first challenges we faced was finding the proper learning rate for the model to learn. For learning rates smaller than 1e-5, the model tended to get stuck with no major decreases in loss after one epoch. On the other hand, when the learning rate was larger than 0.001, the loss function diverged, and the AUC jumped around unstably. After several iterations, we found that the best learning rate was 0.001. We also tried implementing a decaying learning rate approach, which should help the model achieve a better minimum once it is closer to the the optimum. However, this approach did not end up outperforming the best learning rate with an adaptive momentum based optimizer. As for the initialization, although Gaussian initialization worked well, Xavier initalization helped get the model achieve better performance earlier (AUC 0.94 after 1 epoch vs 0.89). However, we found that the final AUC achieved by Xavier initialization was comparable to the Gaussian initialization at the end of training. Surprisingly, we found that He initialization diverged and did not outperform the Gaussian initialization as expected. This initialization, originally derived to deal with a novel Parametric Rectifier Linear Unit (PReLU) \cite{bib11}, perhaps may not apply as well to the LeakyReLU units (a specific case of the proposed PReLU) we ended up using. For data augmentation, we determined the acceptable limits of variation from the original image by manually visualizing several sample outputs. For example, to augment of brightness for each tile, a value was sampled uniformly at random between 0.75 and 1.25; that is, the brightness would not be changed by more than 25\%. Similarly, we chose a maximum contrast difference of $\pm$25\%, saturation difference of $\pm$25\%, and hue difference of $\pm$5\%. We believe that these exact limits could be further tuned, but our heuristic approach already showed a significant boost in performance. Without augmentation, our model was only able to obtain macro AUC around 0.93 on the validation set for the lung cancer task. With data augmentation, we were able to consistently achieve macro AUCs above 0.96. In terms of the optimizers, both Adam and RMSProp performed similarly. SGD, on the other hand, did not perform as well. This is likely because SGD does not have a momentum-type mechanism to adaptively adjust the learning rate. Use of SGD can require a lot more careful hyperparameter tuning than the adaptive methods such as Adam and RMSProp.

